# Expression of the type III secretion system genes in epiphytic *Erwinia amylovora* cells on apple stigmas benefits endophytic infection at the hypanthium

**DOI:** 10.1101/2020.07.23.218156

**Authors:** Zhouqi Cui, Regan B. Huntley, Neil P Schultes, Kaleem U. Kakar, Quan Zeng

## Abstract

*Erwinia amylovora* causes fire blight on rosaceous plants. Flower surfaces are the primary location in the fire blight infection pathway. Here E. amylovora proliferates on stigmatic and hypanthium surfaces as epiphytic growth, followed by subsequent endophytic (intercellular) infection in the hypanthium. The type III secretion system (T3SS) is an important virulence factor in *E. amylovora*. Although the role of T3SS during the endophytic infection is well characterized, its expression during the epiphytic colonization and role in the subsequent infection is less understood. Here, we investigated the T3SS expression in epiphytic *E. amylovora* on stigma and hypanthium of apple flowers, under different relative humidities (RH). On stigma surfaces, T3SS was expressed in a high percentage of E. amylovora cells, and its expression promotes epiphytic growth. On hypanthium surfaces however, T3SS was expressed in fewer *E. amylovora* cells than on the stigma, and displayed no correlation with epiphytic growth, even though T3SS expression is essential for infection. *E. amylovora* cells grown on stigmatic surfaces and then flushed down to the hypanthium displayed a higher level of T3SS expression than cells grown on the hypanthium surface alone. Furthermore, cells pre-cultured on stigma before inoculation on hypanthium caused more infections than cells pre-cultured in a T3SS-repressive medium then used as inoculum, suggesting that the T3SS induction during the stigmatic epiphytic colonization may be beneficial for the subsequent infection. Finally, the epiphytic expression of T3SS is influenced by RH. Higher percentage of stigmatic *E. amylovora* cells expressed T3SS under high RH than under low RH.

## Introduction

Erwinia amylovora is a Gram-negative phytopathogenic bacterium that causes fire blight disease on apple, pear and other rosaceous plants. E. amylovora can infect various parts of the hosts including flowers, fruitlets, shoots, and rootstocks. Among them, infection on flowers, referred to as blossom blight, is of the most importance in disease epidemiology and management (Bubán and Orosz-Kovács, 2003). Flowers provide a nutrient rich and moist environment suitable for the epiphytic growth of *E. amylovora* (Norelli and Brandl, 2004). Flowers also contain natural openings that serve as entry points of *E. amylovora* into the hypanthium interior that normally develops into the apple fruit pulp. Infected flowers quickly wilt and are unable to produce fruits, resulting in yield losses. Moreover, after the initial infection of flowers, pathogens can quickly spread to other parts of the tree, resulting in systemic infection. Annual losses to fire blight and cost of control are estimated over 100 million dollars in the United States alone (Norelli et al., 2003).

*E. amylovora* utilizes several virulence factors to successfully cause disease. Arguably the most essential virulence factor is the type III secretion system (T3SS). The T3SS is a needle-like structure that translocates effector proteins from the pathogen cells into host cells (Notti and Stebbins, 2016). The T3 effectors suppress host immunity and induce plant cell death (Boureau et al., 2006). In *E. amylovora*, the T3SS is encoded by three clusters of 34 hrp, hrc, and dsp genes that encode the secretion apparatus, effectors, and other translocated proteins (Sebaihia et al., 2010). Because of its importance in pathogenicity, regulation of the T3SS in *E. amylovora* has been intensively studied. Many different types of regulators, such as sigma factors (Ancona et al., 2014), two component signal transduction systems (Li et al., 2014), protease (Lee et al., 2018), regulatory small RNAs (Zeng et al., 2013; Zeng and Sundin, 2014; Schachterle et al., 2019), as well as bacterial secondary messengers (Edmunds et al., 2013; Ancona et al., 2015) have been identified to control the expression of the T3SS genes. The underlying yet unproven hypothesis from these studies is that the fire blight pathogen *E. amylovora* senses environmental signals and adjusts T3SS genes expression accordingly.

Bacterial pathogen modulation of virulence gene expression in response to host and environmental signals has been documented (Pusey, 2000; Cui et al., 2015). While virulence factors, such as T3SS, are non-essential to cellular metabolism and growth, their synthesis and assembly are costly in terms of energy and material and can impose a penalty on bacterial growth (Cui et al., 2019). For this reason, pathogens tend to express the virulence genes only when necessary. For instance, in necrotrophic plant pathogen *Dickeya dadantii*, T3SS is highly expressed at early stage of infection but is subsequently diminished at a later stage of infection (Cui et al., 2018b). In *E. amylovora*, expression of key type III secretion genes (*hrp/dspA/E*) were activated in a narrow time window peaking between 24–48 hours post inoculation (Pester et al., 2012). Most work investigating virulence gene expression focused on the endophytic stages of infection, when pathogens are present in the intercellular space of the plant and is directly associated with symptom development, pathogen proliferation.

Although endophytic stage of infection, referring to when pathogens are present in the intercellular space of the plant, drew most research attention as it is directly associated with symptom development, pathogen proliferation and migration epiphytically on surface of plants, as observed in *Pseudomonas syringae* (Yu et al., 2013), *Xanthomonas axonopodis* (Gent et al., 2005) and *Pantoea ananatis* (Coutinho and Venter, 2009), is often an indispensable phase of infection as it prepares pathogens to be ready to invade the host when the environment becomes inductive. Prior to causing infection at the floral cup, *E. amylovora* first colonizes on the stigma surfaces and multiplies epiphytically using the nutrients from the stigma exudates (Pusey, 1997; Pusey et al., 2008). With the aid of free-moving water such as dew and rain, the stigmatic population of *E. amylovora* migrates down to the hypanthium, and initiates the endophytic stage of infection through natural openings called nectarthodes (Thomson, 1986a; Bubán and Orosz-Kovács, 2003). The epiphytic growth of *E. amylovora* is heavily influenced by many environmental factors, such as temperature, relative humidity, and precipitation (Pusey, 2000; Pusey and Curry, 2004). However, whether expression of virulence genes, such as T3SS, is needed during the epiphytic stage of infection, and its impact to the endophytic stage infection is generally less understood.

In this study, we hypothesize that the fire blight pathogen *E. amylovora* grown epiphytically on flowers express T3SS genes according to the spatial, temporal, and environmental signals. We also hypothesize that the variations in T3SS expression is not random, but confers biological significance in promoting pathogen colonization and infection. We monitored the T3SS expression levels at stigma and hypanthium of apple flowers, under different relative humidities (60%, 80%, and 100%), over a period of 3 days. Epiphytic pathogen population size and infection rate of the ovary tissue were also determined in tandem with the T3SS expression to establish any correlation and thereby link T3SS function. Our findings suggest that epiphytic colonization of *E. amylovora* on flowers not only builds up a sizable pathogen population it also induces pathogen virulence, both of which are important for the successful infection at the hypanthium.

## Materials and Methods

### Bacterial strains, plasmids, media

Strains of *E. amylovora* and *Escherichia coli* were cultured in lysogeny broth (LB) medium at 28°C and at 37°C, respectively. All strains were stored at −80°C in 15% glycerol. For *in vitro* induction of T3SS, an overnight culture of *E. amylovora* was sub-cultured in a *hrp*-inducing minimal medium (Yang et al., 2007). When necessary, antibiotics were added to the medium at the following concentrations: chloramphenicol, 30 μg/mL; kanamycin, 50 μg/mL; ampicillin, 100 μg/mL; spectinomycin, 50 μg/mL.

### Construction of dual fluorescence reporter plasmids

Dual fluorescence reporter pP*hrpA-*P*nptII* plasmid was modified from a previously constructed plasmid pAT-*PnptII-gfp-PhrpA3937-mCherry* (Cui et al., 2018b). Briefly, the *hrpA* promoter was PCR amplified from the *E. amylovora* 1189 chromosome, digested as a *SacI* and *BamHI* fragment to replace the promoter *hrpA3937,* resulting in the pAT-P*hrpA*_*Ea*_-P*nptII* plasmid. Then the whole fragment of P*nptII-gfp-PhrpA*_*Ea*_-*mCherry* was PCR amplified using pAT-P*hrpA*_*Ea*_-P*nptII* plasmid as the template, and was digested by *Kpn*I and *Sal*I. The released fragment was subsequently cloned into vector pCL1920 between the same restriction enzyme sites to obtain pP*hrpA-*P*nptII.* The pP*gapA-*P*nptII*, pP*hrpA-*P*hrpJ* and pP*hrpA-*P*dspE* plasmids were subsequently constructed by replacing the *hrpA*_*Ea*_ promoter with the promoter of *gapA*, and by replacing the *nptII* promoter with the promoter of *hrpJ* or *dspE*, respectively. All constructs were confirmed by sequencing and the primer sequences were listed in Table S1.

### Construction of *hrpL* deletion and overexpression strains

The Δ*hrpL* deletion mutant was constructed in a previous study (McNally et al., 2012). Chromosomal *hrpL* overexpression strain of *E. amylovora* was constructed using a previously reported λ red recombinase method to replace the original promoter of *hrpL* with an *nptII* promoter (Datsenko and Wanner, 2000b; Zhao et al., 2009). Briefly, the *nptII* promoter was PCR amplified from plasmid pKD4 (Datsenko and Wanner, 2000a) using primers containing 50-nucleotide homology arms of *hrpL* promoter regions flanking a constitutively expressed promoter *nptII.* The kanamycin resistance cassette gene *kan* was also amplified from plasmid pKD4. The amplified PCR fragments of *PnptII* and *kan* were digested by *EcoR*I / *BamH*I and *Kpn*I / *EcoR*I, respectively. The digested fragments were cloned into vector pBSK+ (Stratagene, La Jolla, CA) to acquire pBSK-Km*-PnptII*. The PCR products of Km*-PnptII* were amplified, purified and electroporated into *E. amylovora* 1189 strain expressing the recombinase gene from the helper plasmid pKD46. The P*nptII-hrpL* strain was selected on LB medium amended with Kanamycin and ampicillin and confirmed by PCR and sequencing.

### Detached apple flower assay

Apple flower clusters were collected from ‘Macoun’ apple trees during bloom, placed on ice, and transferred to laboratory for immediate processing. Freshly opened flowers with petals expanded 80% were detached from flower clusters, placed on 10ml plastic tubes with the petiole submerged in 10% sucrose solution as described by Pusey (Pusey, 2000). Tubes holding the flowers were placed on plastic tube racks enclosed in 4-liter plastic containers with 1 litter glycerol-water mixture for the purpose of maintaining various levels of relative humidity. The correlation curve linking glycerol concentration to relative humidity was established in previous study (Johnson, 1940). The relative humidity was maintained at 60% (62.7% glycerol), 80% (37.5% glycerol) and 100% (no glycerol).

To prepare inoculum, *E. amylovora* 1189 strains were grown overnight in LB broth (adjusting OD_600_=1.0). The overnight culture was collected by centrifugation at 9,000 g for 1 min and resuspended in 0.5 × phosphate buffer saline (PBS) for inoculation. For stigma inoculation, 2 μl of inoculum was evenly distributed among five stigmas on each flower using a micropipette. For hypanthium inoculation, 1 μl of inoculum suspension was directly placed on the hypanthium surface on each flower. For water-flushed treatments, stigma was originally inoculated with *Ea* as described above. Two days post inoculation, 5 μl of sterile water was pipetted onto the stigmas of each flower with gentle mixing, followed by pipetting the water containing *Ea* cells directly to the hypanthium surface. Flowers for the flushing assay were maintained at 80% relative humidity prior to visualizing the expression of *hrpA* at hypanthium 24 and 48 hours after the water flush.

### Shoot and fruitlet inoculation

For shoot inoculation, freshly grown succulent apple shoots (cultivar ‘Gala’) were inoculated with *E. amylovora* by cutting the tip of the youngest shoots with a scissors dipped in a suspension of *E. amylovora* 1189 at 10^8^ CFU/ml. Inoculated shoots were bagged with a plastic Ziploc bags to maintain high humidity conditions. For fruitlets inoculation, fruitlets with approximately 1.5 inches by diameter were sterilized with 10% bleach, wounded by a 0.2-mm syringe needle. Five microliters of an *E. amylovora* 1189 suspension at 10^7^ CFU/ml were placed at each wound. The inoculated fruitlets were incubated at 28°C in an enclosed container with wet paper towels.

### Confocal and epifluorescence microscopy

Fresh stigma, hypanthium, or other plant samples were dissected into 2 mm^2^ sections and mounted on microscope slides, covered by cover slips. The cover slip is secured to the slides with scotch tape. The green and red fluorescence was observed using either a Leica SP5 confocal microscope or a Zeiss Axioplan 2 fluorescence microscope. The Leica SP5 confocal microscope (Leica, Wetzlar, Germany) has four laser channels, 405 nm, multi-line Argon, 561 nM and 633 nm, and two HyD detectors. The Zeiss Axioplan 2 fluorescence microscope (Zeiss, Oberkochen, Germany) contains three fluorescence filter sets: 02 EX G365, 10 EX BP450-490, and 15 EX BP 546/12. Overlay of green and red fluorescence images was performed using Leica LAS-X software (Schneider et al., 2012).

### Quantification of epiphytic *E. amylovora* population

For stigma samples, five stigmas inoculated with *E. amylovora* pP*hrpA-*P*nptII* from each flower were dissected off from the styles and placed in a 1.5 μl microcentrifuge tubes containing 200 μl of 0.5 x PBS. For hypanthium samples, the hypanthium tissue was obtained by removing the corolla, calyx and pedicel. Both liquid present on the hypanthium surface (nectar) and the hypanthium tissue of each flower were collected as a single sample. Hypanthium samples were collected into sterile 1.5 ml Eppendorf tubes with 200 μl of 0.5 x PBS. The tubes were then sonicated for 2 min in a water bath sonicator, followed by vortexing for 15 s. The bacteria-containing PBS buffer was serial-diluted and plated on LB agar plates supplemented with ampicillin for determination cell counts.

### The CyaA translocation assay

The *E. amylovora* full-length DspE-CyaA reporter (DspE_1-737_-CyaA) and the N-terminal DspE fragment-CyaA reporter (DspE_1-15_-CyaA) were constructed in a previous study (Triplett et al., 2009). Overnight cultures of the above strains were adjusted to 1 × 10^7^ CFU/ml in 0.5 x PBS and were sprayed on flowers of apple cultivar ‘Macoun’ of the same stage of bloom. Two days post-inoculation, the stigma portion of each flower was dissected and collected into a 1.5 microcentrifuge tubes. Stigmas of eight flowers were pooled together as one sample. Collected samples were kept on ice during transportation and were stored at −80°C until cAMP determination. cAMP was extracted according to the method reported by Castiblanco et al. (Castiblanco et al., 2018). Briefly, frozen flower stigmas were ground and resuspended in 250 μl of 0.1 M HCl. Suspensions were centrifuge at 3,000 g for 5 min and the cAMP levels in supernatants were measured using the cAMP EIA Kit (Cayman Chemical, Ann Arbor, MI, US) according to the manufacturer’s instructions. Statistical analyses were performed using a one-way ANOVA LSD-test.

### Quantification of disease incidence using inoculum collected from stigma and from an overnight culture in a nutrient rich medium

To prepare inoculum from stigma, overnight cultured *E. amylovora* cells at the concentration of 10^7^ CFU/ml were spray inoculated to open flowers of apple trees (*Malus x domestica* cultivar ‘McIntosh’) maintained at Lockwood farm of the Connecticut Agricultural Experiment Station. The stigma portions of approximately 300 inoculated flowers were harvested two days post-inoculation, and pooled together into a sterile 15 ml centrifuge tube. Ten milliliters of 0.5 X PBS was added into the tube containing the stigmas, which was sonicated for 5 minutes and vortexed for 30 seconds. To prepare inoculum from a nutrient rich, *hrp-*repressing condition, *E. amylovora* 110 was cultured in LB broth overnight at 28°C. Cell counts of live *E. amylovora* from both stigma and LB culture were determined by serial-dilution and plating on LB agar plates.

The inoculum from stigma and from LB overnight culture was serial-diluted and inoculated onto hypanthium of flowers on 6 McIntosh trees. A total of six inoculum treatments, including three serial diluted stigma samples and three serial-diluted LB samples, were inoculated on 18 branches, with each treatment inoculated on three branches. Each branch contains more than 60 individual flowers at stage of bloom. Ten microliters of inoculum was added to the hypanthium by hand with a pipette.

Three weeks after pathogen inoculation, inoculated flowers were rated for the blossom blight symptoms (black withering, dying of the apple flowers, and emergence of ooze droplets). Percentage of diseased individual flowers in the total number of individual flowers was determined for each treatment.

## Results

### *E. amylovora* expresses T3SS genes in a proportion of the total cells

To monitor the expression of T3SS genes *in planta*, we constructed a dual fluorescence reporter plasmid pP*hrpA-PnptII*, which contains an *nptII* promoter-*gfp* transcriptional fusion and an *hrpA* promoter-*mCherry* transcriptional fusion on one plasmid. This construct enables all *E. amylovora* cells to be green fluorescent, while only cells that expressed *hrpA*, a gene encoding the T3 pilus, to be red fluorescent. We observed that among all *E. amylovora* cells under the same *in planta* conditions (shoots, fruitlets, and flowers), only a subpopulation highly expressed *hrpA* while the rest cells did not express *hrpA* (Fig. S1A). The *hrpA-*expressing cells ranged from 35-70% in different conditions tested (Table 1). As a negative control, a non-T3SS gene *gapA*, which encodes a glyceraldehyde-3-phosphate dehydrogenase, was evenly expressed in all cells (Fig. S1B).

**Table 1:**
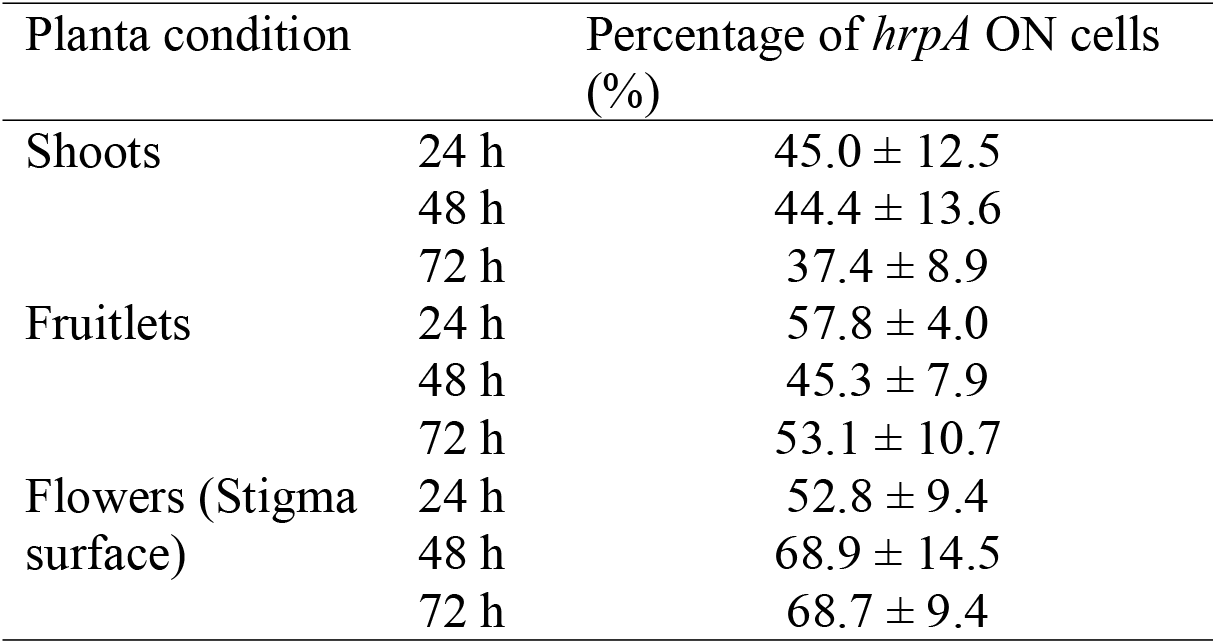
Percentage of *hrpA* expressing cells in total *E. amylovora* population under *in planta* colonization.

To determine if cells that highly expressed *hrpA* also highly expressed other T3SS genes, we constructed two other dual reporters pP*hrpA-*P*dspE* and pP*hrpA-*P*hrpJ*, and observed the co-expression of *hrpA* with *dspE* (encoding a T3 effector) or *hrpJ* (encoding a T3 secreted protein). When cultured in the *in vitro hrp*-inducing medium, cells highly expressing *hrpA* also simultaneously highly expressed *dspE* or *hrpJ* (Fig. 1). Taken together, these observations suggests the expression of T3SS genes occurs in an organized, tandem manner to allow the functional establishment of T3SS in a sub-population of *E. amylovora* cells. By adjusting the percentage of “ON” cells in a population, *E. amylovora* adjust the overall expression level of T3SS similar to what was reported in other pathogen systems (Cui et al., 2018a). Hereafter, we used the percentage of the *hrpA-*expressing cells to represent the percentage of T3SS-expressing cells and the overall T3SS expression in a population.

**Figure 1.**
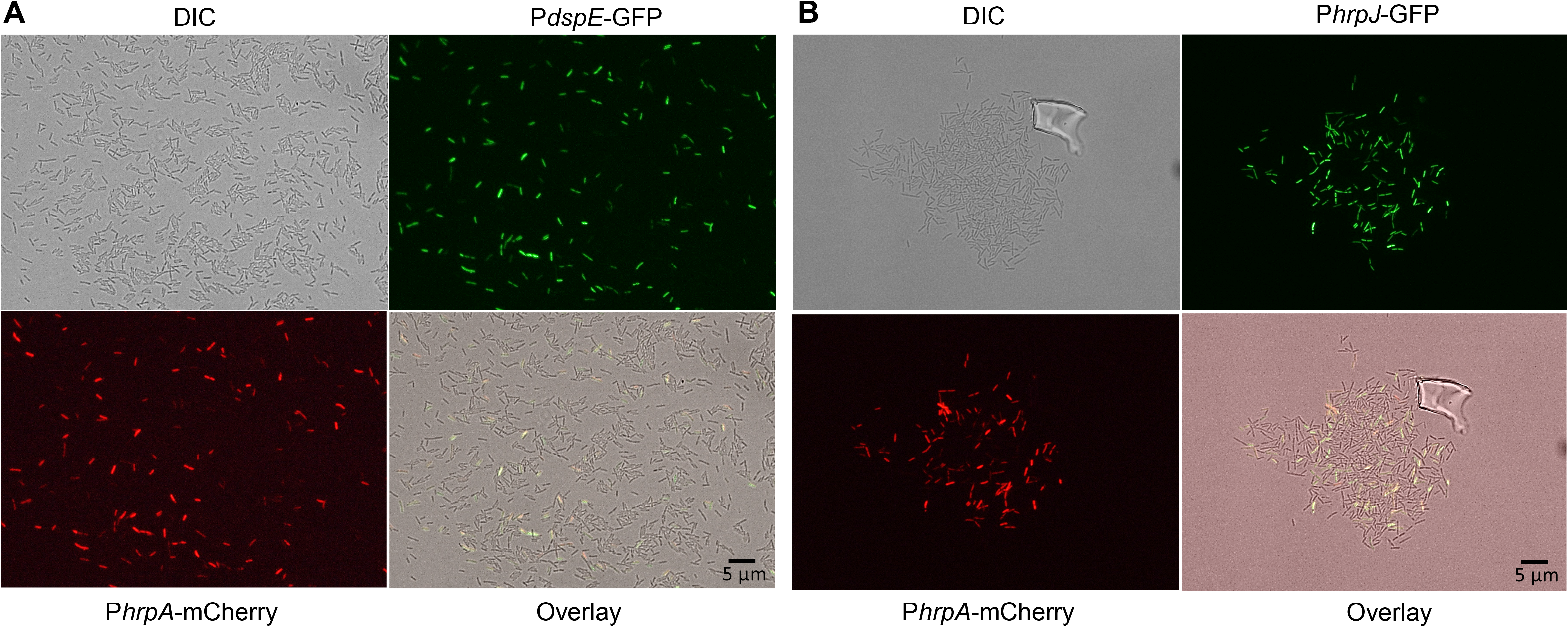
Fluorescence microscopy observation of *E. amylovora* harboring (A) pP*hrpA-*P*dspE* and (B) pP*hrpA-*P*hrpJ* when cultured in Hrp-inducing minimal medium. *E. amylovora* 1189 carrying the dual-fluorescence promoter reporter plasmid was first cultured overnight in LB. Cells from the overnight culture were collected by centrifugation and transferred to the Hrp-inducing minimal medium for 18 h. Differential interference contrast (DIC) light-microscopic observation (upper left panel), along with green fluorescence (upper right panel) and red fluorescence (lower left panel) were observed in the same group of cells. The lower right panel is the overlay of the green and red fluorescence images.

### On stigma, T3SS expression promotes the epiphytic growth of *E. amylovora* and is positively impacted by relative humidity

On stigma surfaces, the percentage of *hrpA-*expressing *E. amylovora* cells increased as the colonization progressed (Fig. 2A). By Day 3, 70% of total cells expressed *hrpA* (Fig. 2A). We observed that as the relative humidity rose the percentage of *E. amylovora* cells in a population that expressed *hrpA* also increased (Fig. 2A &D), suggesting that relative humidity positively impacts the expression of T3SS in *E. amylovora* during its colonization on stigma. Interestingly, relative humidity also had an impact on the growth of *E. amylovora*. Higher relative humidity resulted in a higher epiphytic population of *E. amylovora* on stigma than the populations grown at lower relative humidites (Fig. 2B). A positive correlation between T3SS expression and bacterial growth was observed (R^2^=0.98; Fig. 2C). These observations support the idea that the presence of T3SS benefits epiphytic growth of *E. amylovora* on stigmas.

**Figure 2.**
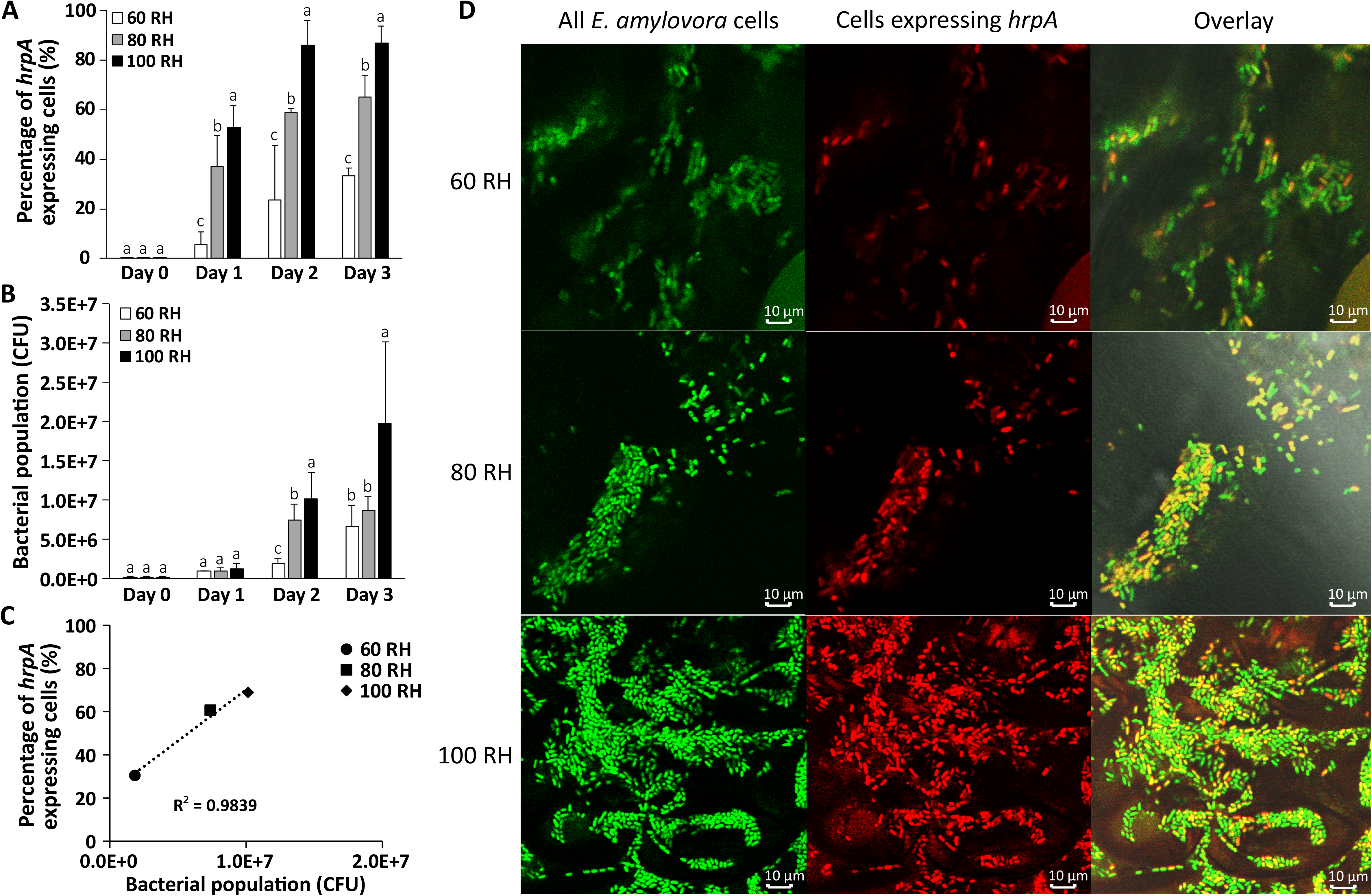
Expression of *hrpA* in epiphytic *E. amylovora* cells on apple stigma surfaces and its correlation with bacterial growth under various relative humidities (RH). **(A)** Percentage of *hrpA*-expressing cells (*hrpA* ^ON^ cells) in the total *E. amylovora* cells on stigma surfaces when flowers were maintained under different RH (60%, 80% and 100%) within the first three days after inoculation. *E. amylovora* carrying the dual reporter pP*hrpA-*P*nptII* was inoculated onto the stigma protion of detached apple flowers. The percentage was obtained from the number of green and red fluorescent bacterial cells observed with a fluorescence microscope. At least five images were taken at each time point. Different letters denote statistical significance (*P* < 0.05, identified by ANOVA). **(B)** *E. amylovora* cell counts on stigmas under 60%, 80% and 100% RH within the first three days post-inoculation. Flowers used for cell count determination were maintained under the same experimental setting as the flowers used for the *hrpA* expression monitoring. The cell counts were determined by plating on LB agar plates. **(C)** Correlation between the percentage of *hrpA* ^ON^ cells and the bacterial population under 60%, 80% and 100% RH at 48 hpi. **(D)** Visualization of *hrpA* ^ON^ cells in total *E. amylovora* cells on stigma under 60%, 80% and 100% RH two days post inoculation using a confocal microscope. All *E. amylovora* cells were visualized by green fluorescence and the *hrpA* ^ON^ cells were visualized by red fluorescence.

To test this hypothesis, we compared the epiphytic growth of the wild type *E. amylovora*, a T3SS mutant (Δ*hrpL*), and an *E. amylovora* strain overexpressing *hrpL* (*nptII-hrpL*) on the stigma surface. A clear growth retardation was observed when T3SS was inactivated (Δ*hrpL*), while overexpression of T3SS resulted in slightly faster growth than the wide type (Fig. 3A). This data, in combination with the observation that the epiphytic growth of *E. amylovora* and its T3SS expression are positively correlated, together suggests that the expression of T3SS supports epiphytic growth of *E. amylovora* on stigmatic surfaces.

**Figure 3.**
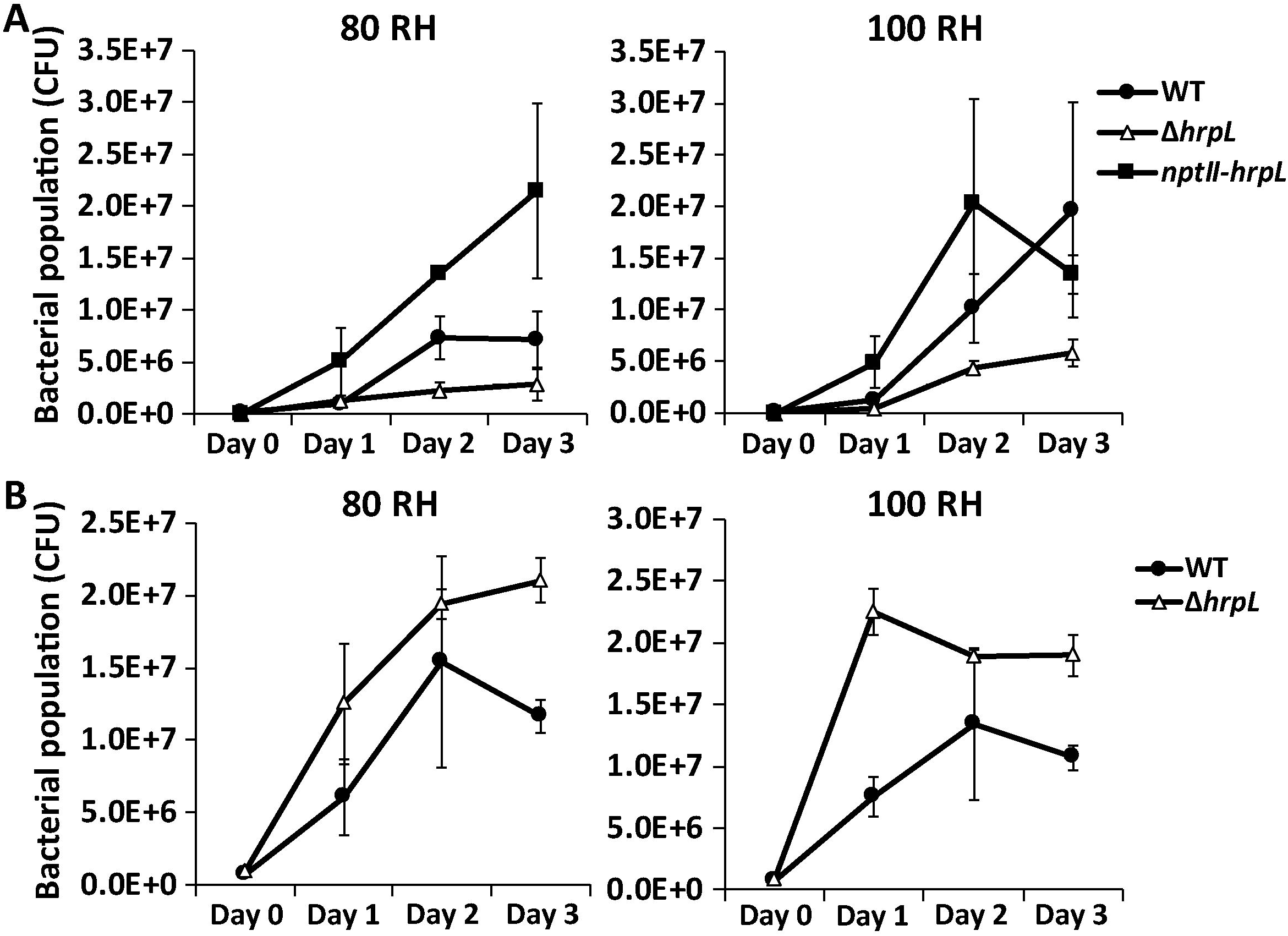
Growth of *E. amylovora* on stigma (A) and hypanthium (B) surfaces in the presence and absence of T3SS. *E. amylovora* wild type (WT) and *hrpL* deletion mutant (Δ*hrpL*) were inoculated onto stigma (**A**) and hypanthium (**B**) surfaces of detached apple flowers. The inoculated flowers were maintained under 80% and 100% RH. Cell counts were determined by plating on Day 0 to Day 3. *E. amylovora* carrying an overexpression construct of *hrpL* (*nptII-hrpL*) was also tested on stigma. Data points and error bars represent the means and standard deviations of three biological replicates.

### DspE is translocated into host cells on stigma

To determine if the expression of T3SS genes in stigma-grown *E. amylovora* cells leads to the translocation of T3 effectors into host cells, we used a previously established translocation assay with a T3 secreted effector DspE fused with an adenylate cyclase (CyaA) gene (Triplett et al., 2009). Elevated cyclic AMP levels were detected in stigmas inoculated with *E. amylovora* carrying the DspE_(1-737)_-CyaA reporter, compared to stigmas from non-inoculated flowers or inoculated with *E. amylovora* carrying a DspE with a truncated translocation signal (DspE_(1-15)_-CyaA) (Fig.4). This confirmed that T3SS is not only actively expressed on stigma, but actively functions as a protein translocation mechanism delivering effector proteins into host cells.

**Figure 4.**
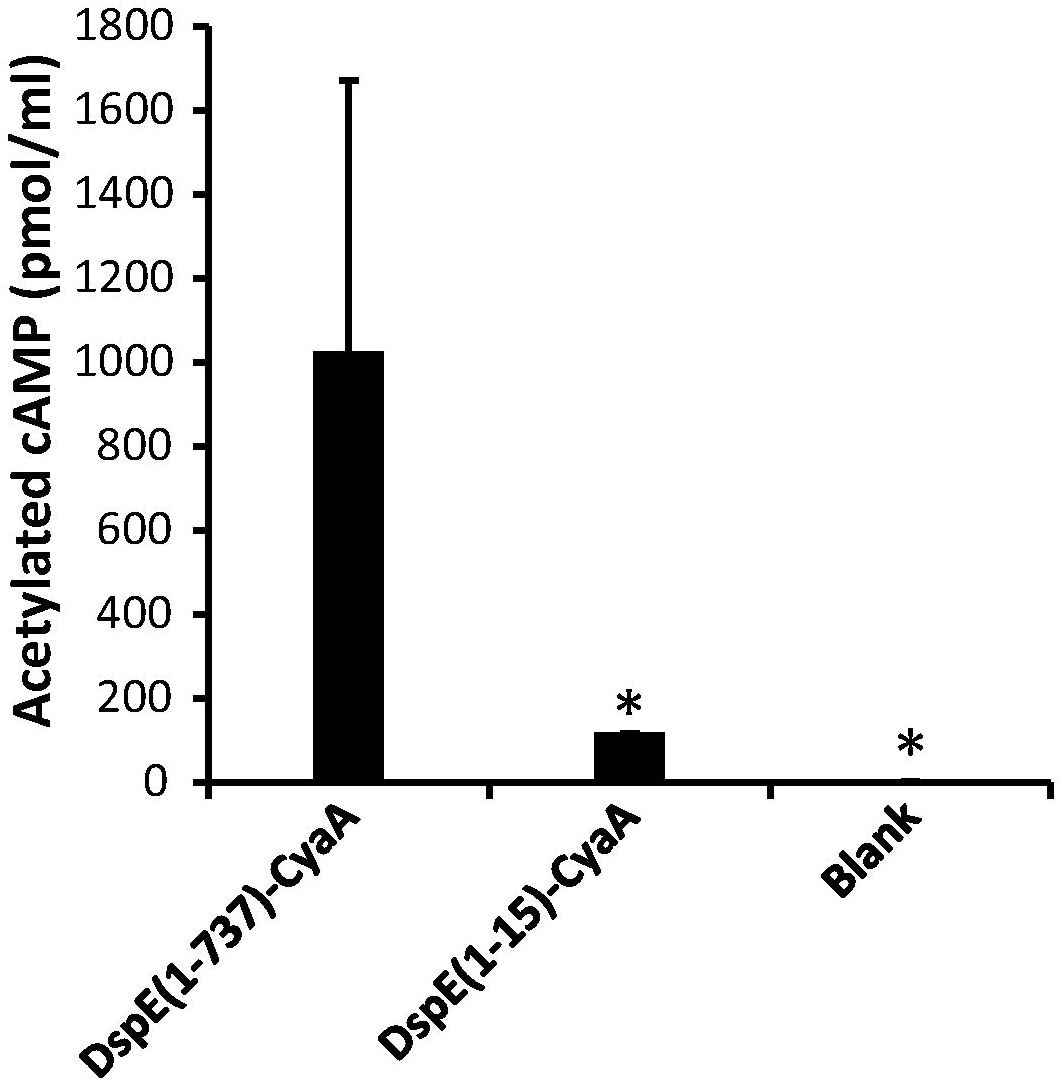
Translocation of T3 effector DspE into stigma cells. cAMP accumulation was determined in apple stigmas inoculated with *E. amylovora* 1189 expressing DspE_(1-737)_-CyaA and DspE_(1-15)_-CyaA. Each sample contained stigmas of eight flowers. Un-inoculated flowers were used as blank control. Data points and error bars represent the means and standard deviations of three biological replicates. Stars denote statistical significance (*P* < 0.05, identified by ANOVA).

### T3SS expression in *E. amylovora* on the hypanthium surface is much lower than on the stigma, is not related to epiphytic growth, but is critical for infection

Compared to on stigma surfaces, expression of T3SS in epiphytic *E. amylovora* cells is much lower at the hypanthium surfaces (Fig. 5, Fig. S2). On Day 1, almost no cells expressed *hrpA* on hypanthium surfaces. The percentage of *hrpA*-expressing cells barely reached 25% by Day 3. The fact that T3SS expression is very low, especially at the initial stage of colonization, suggests T3SS may not be beneficial for the epiphytic growth of *E. amylovora* on the hypanthium surface as it is on the stigma. To test this hypothesis, we compared the epiphytic growth rate of the wild type *E. amylovora* with growth rate of the Δ*hrpL* on hypanthium surface. We did not observe any growth retardation in Δ*hrpL* in comparison with the wild type (Fig. 3B), which confirmed that T3SS expression is not beneficial for *E. amylovora* to colonize the hypanthium surface as it is on the stigma.

**Figure 5.**
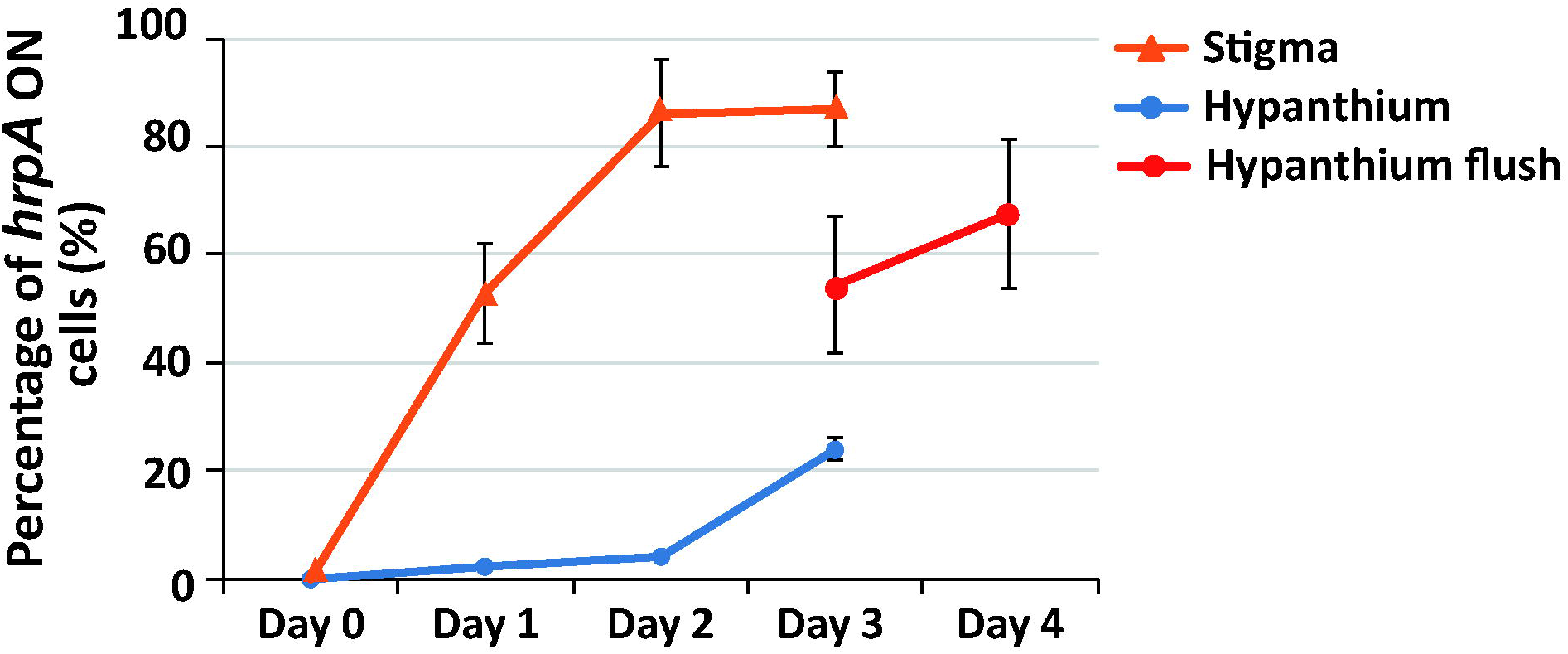
Percentage of *hrpA* ^ON^ cells in the total population on stigma surfaces (orange line), hypanthium surfaces (blue line), and on hypanthium surfaces with inoculum acquired from stigma surfaces through water flushing (red line). *E. amylovora* carrying the pP*hrpA-*P*nptII* dual reporter were inoculated directly onto stigma surface (blue line), hypanthium surface (orange line), and first on stigma surface and allowed to grow for two days and flushed down to the hypanthium with 5 μl of water on Day 2 (red line). The flower samples were maintained under 80% RH and their green and red fluorescence was determined using a confocal microscope. Data points and error bars represent the means and standard deviations of three biological replicates.

The hypanthium surface is not only nutrient rich and able to support epiphytic growth of microbes, but it also contains natural openings (nectarthodes) where pathogens enter into the hypanthium and cause infection. To determine whether T3SS is needed for *E. amylovora* to cause infection in the hypanthium, next we inoculated the wild type and Δ*hrpL E. amylovora* on the hypanthium surface and evaluated the blossom blight symptoms at a later time (3-4 days). Interestingly, even though Δ*hrpL* grew as well as the wild type on the hypanthium, flowers inoculated with Δ*hrpL* remained healthy whereas 92% of flowers inoculated with the wild type developed water soaking and blight symptoms three days and four days after inoculation, respectively (Fig. 6).

**Figure 6.**
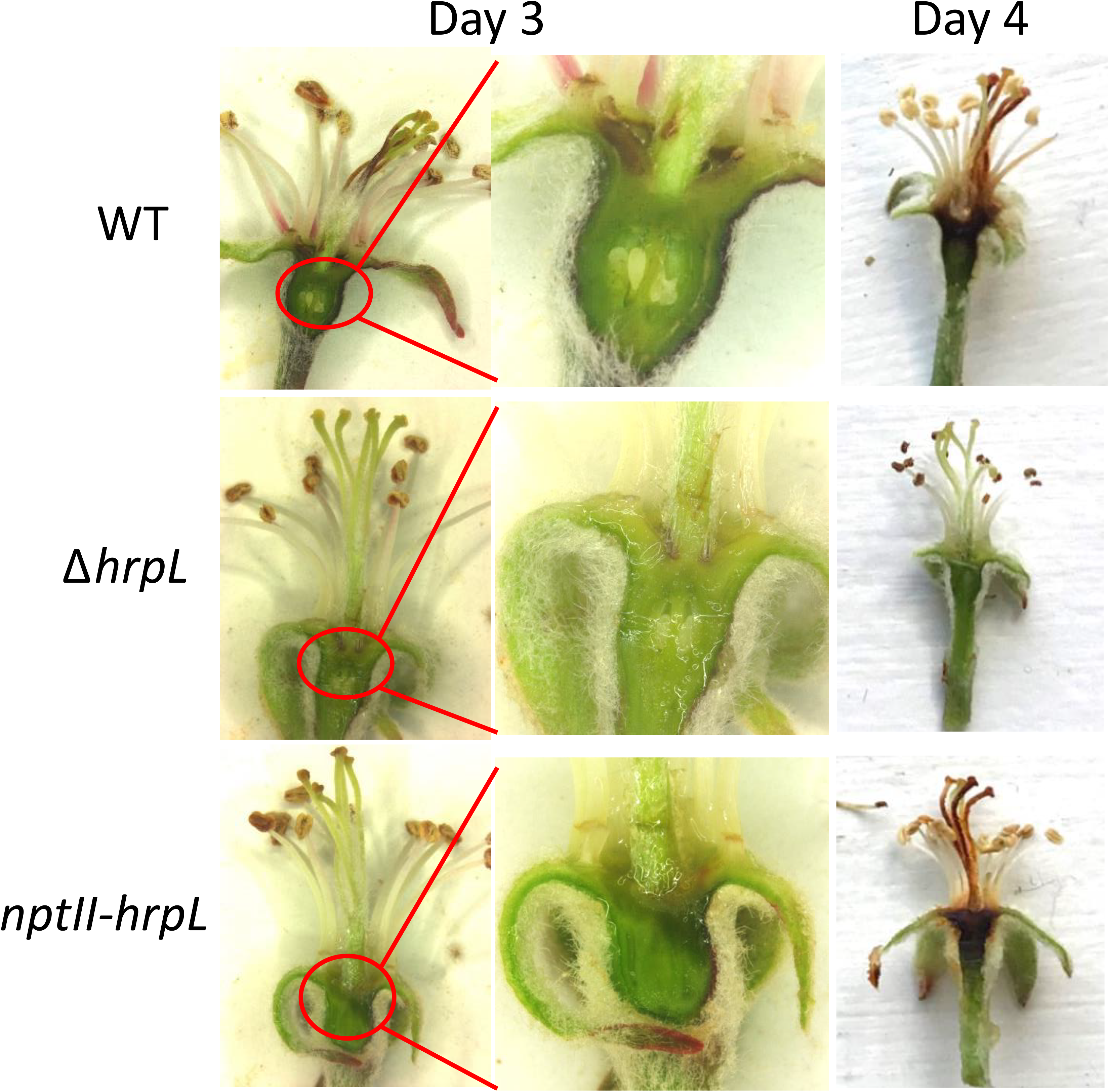
Disease symptoms caused by *E. amylovora* wild type (WT), *hrpL* deletion mutant (Δ*hrpL*) and *E. amylovora* with an overexpression construct of *hrpL* (*nptII-hrpL*) when inoculated on hypanthium surface after three and four days. *E. amylovora* at the concentration of 10^8^ CFU/ml was inoculated onto the hypanthium surfaces. Inoculated flowers were maintained at 28°C and at 100% RH prior to disease symptom examination.

If T3SS expression is indeed needed for infection in the hypanthium, we would expect that the level of T3SS expression would be higher in endophytic *E. amylovora* cells present internally in the infected hypanthium tissue, even though T3SS expression levels are low for hypanthium surface grown cells. Indeed, a much higher level of *hrpA* expression was observed in the internal tissue of the hypanthium (~60%, Fig. 7) than that on the hypanthium surface (< 25%). Taken together, our data suggests that T3SS expression is not needed for the epiphytic growth of *E. amylovora* at the hypanthium surface, but is required for the subsequent endophytic infection inside the hypanthium.

**Figure 7.**
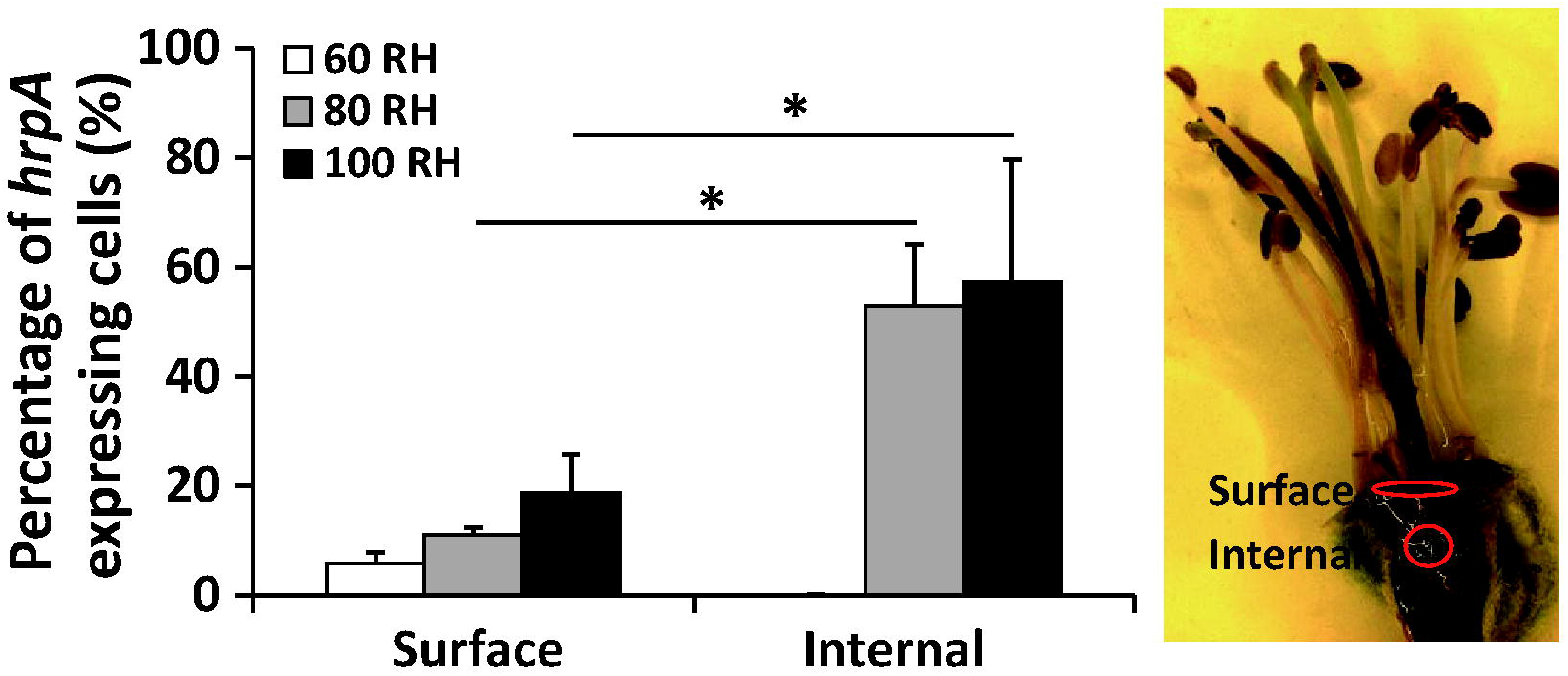
Expression of *hrpA* in *E. amylovora* grown epiphytically on hypanthium surfaces and endophytically in infected internal tissue of hypanthium at 60, 80, and 100% RH. *E. amylovora* carrying the pP*hrpA-*P*nptII* dual fluorescent reporter was inoculated on hypanthium surfaces and maintained at 60, 80, and 100% RH. Four days post-inoculation, green and red fluorescence of samples collected from hypanthium surfaces and from the infected tissue located internally in the hypanthium was observed using fluorescence microscope. Data points and error bars represent the means and standard deviations of three biological replicates. Stars denote statistical significance (*P* < 0.05, identified by ANOVA).

### Epiphytic growth of *E. amylovora* on stigma primes up its virulence prior to hypanthium infection

The stigma surface is a T3SS-inducing environment, whereas hypanthium surface is a T3SS-repressive environment. We suspect that during natural infection, the colonization of *E. amylovora* on the stigma surface prior to the infection of the hypanthium may not only benefit the occurrence of infection by building up the pathogen population, but also by priming up pathogen’s virulence. To test this hypothesis, we measured the *hrpA* expression in *E. amylovora* cells directly inoculated on the hypanthium and in cells that first grew on the stigma and were then flushed down to the hypanthium. *hrpA* expression is significantly higher in cells flushed from stigma down to the hypanthium (red line) than in cells that had been grown on the hypanthium surface from the beginning (blue line in Fig. 5). This suggests that stigma colonization indeed primes *E. amylovora* virulence prior to reaching to the hypanthium.

### *E. amylovora* cells epiphytically grown on the stigma have a better chance to infect the hypanthium interior than *E. amylovora* cells grown in a *hrp-*repressing, nutrient rich medium

To determine whether the virulence-primed-up *E. amylovora* cells grown on stigma would have a better chance to infect than cells without such virulence induction, we prepared two different inoculums: the first inoculum consisted of virulence-primed *E. amylovora* cells collected from *E. amylovora-*inoculated stigmas and the second inoculum consisted of virulence-suppressed *E. amylovora* cells grown overnight in nutrient rich broth (LB broth, Fig. S3). The two inoculums were serial-diluted and inoculated onto the hypanthium surface to allow blossom blight symptom development. *E. amylovora* cell counts of the each inoculum were also determined by plating. Flowers that received the same inoculum source, but with increasing pathogen numbers resulted in higher infection rates, suggesting pathogen population size is positively correlated to the infection rate (Fig. 8). This was observed with both inoculum types. However, when we compared the infection rate of the two different inoculum sources, we noticed that stigma-reared *E. amylovora* had a higher infection potential (infection/inoculum cell number) than cells cultured in LB broth. Stigma-grown cells required a population size 10^2^-10^3^ times lower than the LB grown cells to achieve a similar rate of infection (Fig. 8). This suggests that enhanced T3SS gene expression in stigma grown epiphytic cells benefits infection later at the hypanthium.

**Figure 8.**
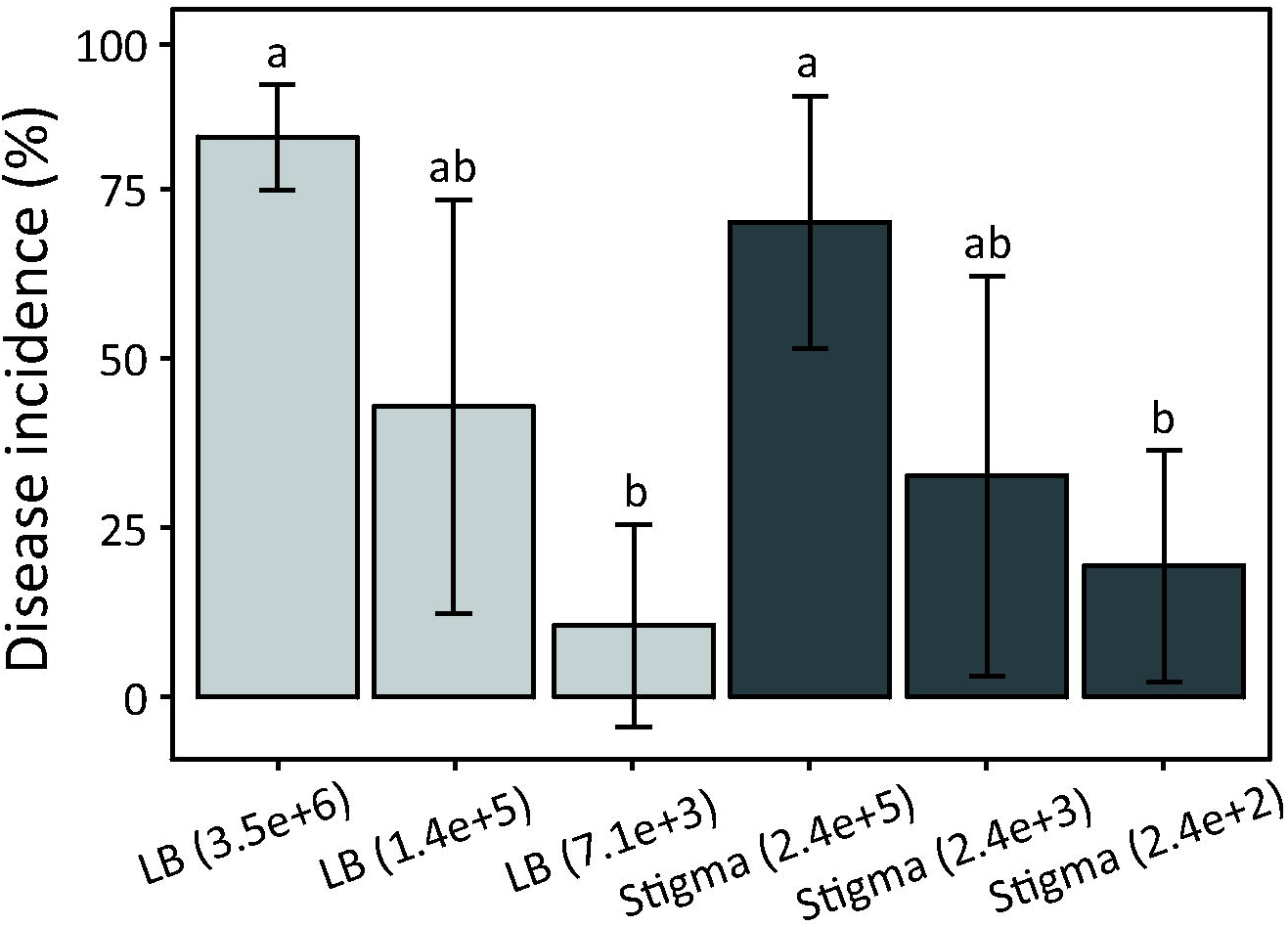
Percentage of apple flowers that developed fire blight symptoms in the field with different inoculums collected from stigma and LB broth. Inoculum from stigma is prepared by first spray-inoculating stigmas of freshly opened flowers with an *E. amylovora* suspension (10^7^ CFU/ml). Stigma portion of the flowers were cut by scissors and collected into a 15 ml centrifuge tube with 0.5 x PBS two days after inoculation. *E. amylovora* cells were washed off from the stigma surfaces by vortexing and sonication. The LB inoculum was prepared by culturing *E. amylovora* in LB broth overnight. Inoculums of the two sources were serial-diluted. A proportion of the inoculum was pipetted to the hypanthium surface of individual flowers, and the rest of the inoculum was plated on LB plates for measuring bacterial cell counts (numbers in brackets). The experiment was performed on apple trees in open field at Lockwood Farm, Hamden, CT. Three replicates were included in each treatment with at least 50 flowers in each replicate. Symptoms of individual flowers were determined three weeks after the original inoculation. Different letters denote statistical significance (*P* < 0.05, identified by ANOVA).

## Discussion

Prior to this study, research determined the major steps in establishing floral fire blight infections. It was understood that *E. amylovora* initially colonizes and propagates mainly on apple stigma establishing a large epiphytic population (Thomson, 1986b; Wilson and Epton, 1989; Johnson et al., 2009a). With the help of free-moving water, such as rain or dew, the pathogen cells migrate from stigma down to the hypanthium and enter the host through nectarthodes (Pusey, 2000). It is also known that the T3SS is a key virulence factor in *E. amylovora* (OH et al., 2005), and expression of *dspE* was observed on stigma (Johnson et al., 2009b). Johnson et al also showed that *hrpL* was also shown to promote epiphytic growth on stigma (Johnson et al., 2009b). However, there is a lack of systemic study understanding the expression of T3SS genes in epiphytic *E. amylovora* present on different flower parts and its biological functions in regards to blossom blight infection. Our study suggests the epiphytic colonization of *E. amylovora* on stigma plays a more important role in the flower infection process than previously thought. First, growth of *E. amylovora* on the stigma surface is much faster than on the hypanthium surface (Fig. 3), most likely due to lower osmolality (resulting from a lower sugar content in stigma exudate) than is present in nectar at the hypanthium. Second, *E. amylovora* colonization on stigma prior to migrating down to the hypanthium primes virulence (by enhancing T3SS expression), thereby increases the infectivity of *E. amylovora* once gaining entry to the hypanthium interior. Stigma-grown *E. amylovora* cells with T3SS expression induced would have enough T3SS structures present for infection, even though they would encounter the T3SS repressive conditions at the hypanthium surface. A summary of the expression dynamics and biological functions of T3SS during the blossom blight infection is illustrated in Fig. 9.

**Figure 9.**
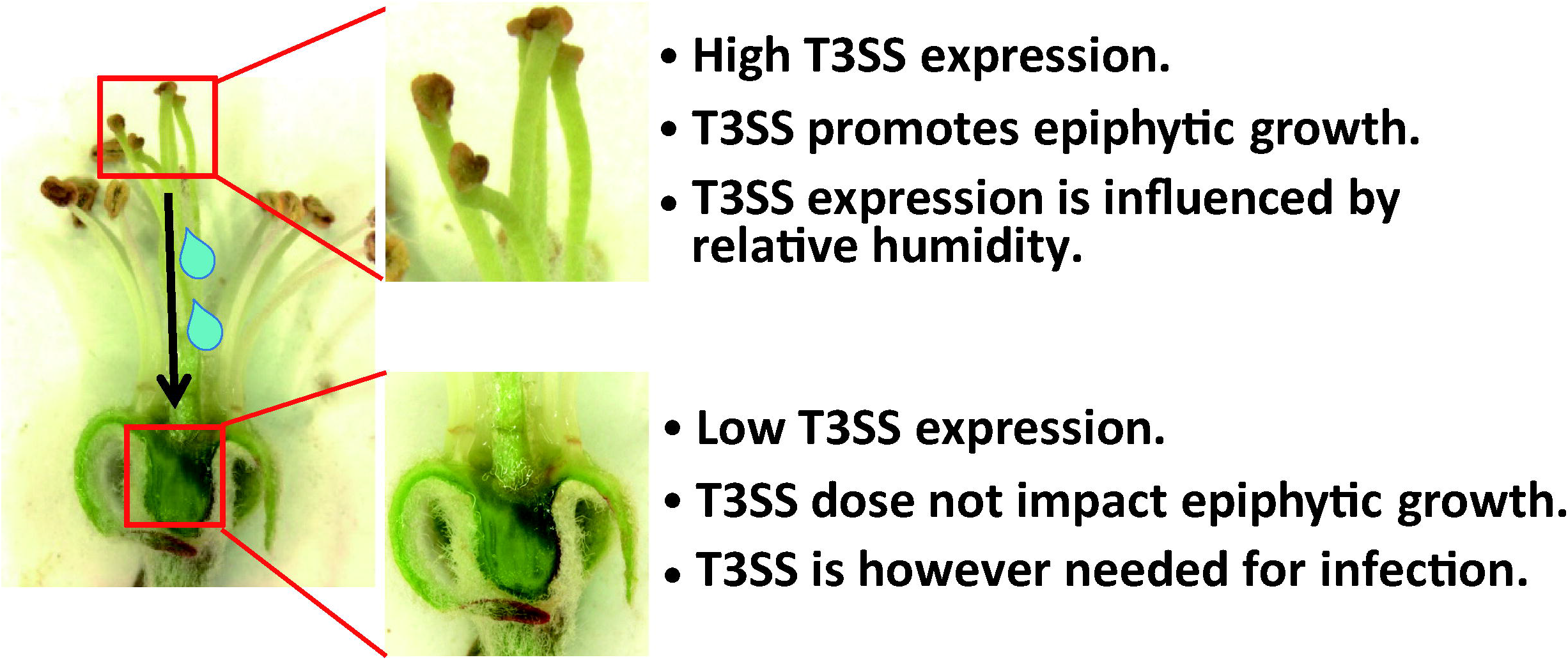
Illustration of the expression dynamics of T3SS genes in *E. amylovora* when colonizing stigma and hypanthium, and its role in epiphytic multiplication of the pathogen and infection. *E. amylovora* was spread to stigma and initially grows epiphytically on stigma surface. On stigma, *E. amylovora* expresses high levels of T3SS genes. The translocation of the T3 effector DspE causes host cell death and helps the bacteria to obtain host resources and build up a large pathogen population. The T3SS expression of *E. amylovora* on stigma is highly influenced by relative humidity: more cells turned on T3SS under high humidity than at low humidity. A wetting event (rain, dew) moves pathogen cells from stigma down to the hypanthium. The hypanthium, possibly due to nectar’s high sugar concentration and the lack of direct host cell contact blocked by the epidermis, represents a T3SS-repressive environment. Epiphytic growth of *E. amylovora* on hypanthium does not require a functional T3SS. However, T3SS function is however needed for the successful infection of the internal tissue of hypanthium. The epiphytic growth of *E. amylovora* on stigma surfaces primes the pathogen’s virulence which helps to overcome the T3SS repressive condition at the hypanthium surfaces and increases its chance of successful invasion at the internal tissue of hypanthium.

An assembly of microorganisms (collectively known as the phytobiome) is known to be able to colonize various nutrient-rich plant surfaces such as roots, leaves and flowers and grow epiphytically. The dominant assumption is that these microbes passively take up the excessive carbohydrates, amino acids, and aromatic organic acids produced and exuded by plants to support the epiphytic growth (Levy et al., 2018). Many plant pathogenic bacteria, such as *Pseudomonas syringae* and *E. amylovora*, also require an epiphytic growth phase prior to subsequent endophytic infection, although the epiphytical stage of colonization is usually asymptomatic. Previous studies of the epiphytic growth of plant pathogens have mainly focused on the stress tolerance aspect (Pfeilmeier et al., 2016), marked by the production of exopolysaccharide (EPS) (Ghods et al., 2015; Prada-Ramirez et al., 2016), and swimming motility as key activities in such process. Predominantly, pathogen virulence gene expression, such as T3SS, has been viewed and investigated during the endophytic phase of infection (Pfeilmeier et al., 2016). Here we showed that expression of T3SS genes may benefit the epiphytic growth of *E. amylovora* during the epiphytic colonization of flowers in a tissue dependent manner: T3SS expression is beneficial for the epiphytic growth on stigma, although not required for the growth of *E. amylovora* at the hypanthium surface where the expression of such genes is kept at low levels. Through the CyaA translocation assay, we demonstrated that T3 effector DspE was indeed translocated into host cells on stigmatic surfaces. DspE, a member of the AvrE superfamily effector, is known to induce host cell death. In fact, browning was often observed on stigmas infected with *E. amylovora*. The induction of host cell death by DspE can likely result in the release of host nutrients thus supporting pathogen’s growth on stigmatic surfaces. This observation, in agreement with a recent report showing that T3SS is beneficial for the epiphytic growth of *P. syringae* on tomato leaves (Stauber et al., 2012), suggests pathogens utilizing plant nutrients on plant surfaces could be a proactive rather than passive process.

Regarding the tissue dependent aspect, we noticed through our observation of the stigma and hypanthium tissue using microscopy, that an obvious layer of epidermis was present on hypanthium surface but not on stigma surface. As the outer epidermal wall, which bears most of the stress exerted by growing internal tissues, is considerably thicker than the inner periclinal and anticlinal walls (Glover, 2000). Having such epidermis present may prevent the T3SS functioning as the protein translocation mechanism at the hypanthium. The sugar-rich nectar at hypanthium may also serve as an environment signal to repress the expression of the T3SS genes.

T3SS genes could be induced by host signals when the pathogens are present in plant apoplast (Yang et al., 2008; Haapalainen et al., 2009). However, whether environmental signals such as relative humidity would also affect the expression of T3SS is not well understood. Fire blight infections occur more often and are more severe under humid conditions or after rain compared to dry weather conditions. However, the correlation of the water to disease severity is often explained as *E. amylovora* grows faster under the humid condition than under the dry condition as shown by Pusey (Pusey, 2000). In this study, we further demonstrated that the humid condition not only favors the growth of *E. amylovora*, but also induces more pathogen cells to express virulence genes (T3SS) than during dryer conditions. As virulence gene expression is critical for infections, enhanced T3SS expression could lead to increased infection rates. This was proven in our inoculation assay using the T3SS induced *E. amylovora* cells pre-cultured on stigma and cells grown in a T3SS repressive condition, that the cells with the T3SS gene expression induced can cause similar level of disease with at least 10-fold less inoculum than cells that are not induced (Fig. 8). Previous studies have shown that pathogen is able to create an aqueous intercellular space in the apoplast endophytically, which benefits the subsequent infection (Xin et al., 2016). Findings from this study highlight the importance of water and relative humidity during the epiphytic colonization and infection of plant pathogenic bacteria. In regard to fire blight management, perhaps the role of water may need to be taken consideration during disease modeling as most fire blight prediction models solely rely on the temperature as the only parameter in evaluating the risk of blossom blight infection.

We observed that even though all *E. amylovora* cells are exposed to the same host environment, the expression of the T3SS genes is restricted to a sub-population of all pathogen cells. This phenomenon was observed in both epiphytic and endophytic stages of infection. This phenomenon was first reported by Johnson et al (Johnson et al., 2009b), although was not further investigated for its functionality and expression pattern. Here, using a dual fluorescent reporter system, we clearly marked the total *E. amylovora* population and the subpopulation that expressed T3SS, thus clearly illustrated the bi-stable nature of the T3SS expression. Furthermore, we showed that different *hrp* and *dsp* genes are simultaneously expressed in the “ON” cells but not in the “OFF” cells. Finally, the percentage of “ON” cells varied at different host environment (e.g. shoots, flowers, fruitlets). These observations support the hypothesis that *E. amylovora* expresses the T3SS genes and establishes the complex protein secretion structure only in a sub-population during infection of the host. It also suggests that *E. amylovora* modulates the overall T3SS expression in a population by adjusting the ratio of the “ON” and “OFF” cells. The incentives of not having the T3SS “turned on” in all bacterial cells may be due to the fact that establishing this none-essential, complex protein secretion system is energy costly and may impose a growth penalty as observed in other plant and animal pathogens (Johnson et al., 2009b; Sturm et al., 2011; Rufián et al., 2016; Cui et al., 2018b; Cui et al., 2019), or perhaps to avoid triggering host immunity.

## Acknowledgments

This study was supported by the Northeastern IPM Center partnership grant, USDA-NIFA-Agricultural Microbiome, USDA-NIFA-Organic Transitions, and USDA-Specialty Crop Block Grant (SCBG) through the Department of Agriculture, State of Connecticut. We thank Jacquelyn La Reau, R. Cecarelli, R. Hannan, and M. McHill for their excellent technical support.

---

**Supplementary Figure S1. Microscopy visualization of *hrpA-*expressing cells at shoots, fruitlets and flowers of apple (A) and *gapA* (a non-T3SS gene)-expressing cells at apple flowers (B) in an isogenic *E. amylovora* population.** *E. amylovora* 1189 carrying either a pP*hrpA-*P*nptII* or a pP*gapA-*P*nptII* dual reporter was inoculated in apple shoots, immature fruitlets and on flower stigmas. Total bacterial cells were visualized by green fluorescence and cells expressed *hrpA* or *gapA* were visualized by red fluorescence.

**Supplementary Figure S2. Fluorescence microscopy observation of the *hrpA* expression in *E. amylovora* on hypanthium surfaces.** *E. amylovora* 1189 carrying a dual fluorescence reporter pP*hrpA-*P*nptII* was inoculated onto apple hypanthium surfaces and observed under a fluorescent microscope within the first three days post inoculation. Flower samples were maintained under 100% RH. All *E. amylovora* cells were visualized by green fluorescence and the *hrpA* expressing cells were visualized by red fluorescence.

**Supplementary Figure S3. Fluorescence microscopy observation of the *hrpA* expression in *E. amylovora* cells cultured in LB broth overnight.** *E. amylovora* 1189 carrying the dual-fluorescence promoter reporter plasmid was cultured in nutrient rich LB broth for 18 h. Differential interference contrast (DIC) light-microscopic observation (left panel), along with green fluorescence (middle panel) and red fluorescence (left panel) were observed in the same group of cells.

