## Supplemental Table 1 for "Expression of the type III secretion system genes in epiphytic *Erwinia amylovora* cells on apple stigmas benefits endophytic infection at the hypanthium"

**Table S1** Bacterial strains, plasmid and primers used in this study.

| Strain or plasmid | Characters and sequences (5’ to 3’)^a^ | Source or reference |
| --- | --- | --- |
| Strains |  |  |
| *Escherichia coli* DH5α |  |  |
| *Erwinia amylovora* |  | [1] |
| Ea1189 | Wild type | [2] |
| *ΔhrpL* | *hrpL* deletion mutant, Cm^r^ | [3] |
| *nptII*-*hrpL* | *hrpL* overexpression mutant, Km^r^ | This study |
| Ea1189 (*nptII*-P*hrpA*) | *E. amylovora* containing dual-fluorescence promoter reporter plasmid *nptII-gfp-hrpA-mCherry* | This study |
| Ea1189 (*nptII*-*gapA*) | *E. amylovora* containing dual-fluorescence promoter reporter plasmid *nptII-gfp-gapA-mCherry* | This study |
| Ea1189 (*dspE*-*hrpA*) | *E. amylovora* containing dual-fluorescence promoter reporter plasmid *dspE-gfp-hrpA-mCherry* | This study |
| Ea1189 (*hrpJ*-*hrpA*) | *E. amylovora* containing dual-fluorescence promoter reporter plasmid *hrpJ-gfp-hrpA-mCherry* | This study |
| *ΔhrpL* (*nptII*-*hrpA*) | *E. amylovora ΔhrpL* containing dual-fluorescence promoter reporter plasmid *nptII-gfp-hrpA-mCherry* | This study |
| *nptII*-*hrpL* (*nptII*-*hrpA*) | *E. amylovora nptII*-*hrpL* containing dual-fluorescence promoter reporter plasmid *nptII-gfp-hrpA-mCherry* | This study |
| Plasmids |  |  |
| pBSK+ |  | Stratagene |
| pKD3 | Cm^r^ , mutagenesis cassette template | [4] |
| pKD4 | Km^r^ , mutagenesis cassette template | [4] |
| pKD46 | Ap^r^ , expresses bacteriophage red recombinase | [4] |
| pAT-P*nptII*-gfp-P*hrpA*3937-mCherry | pPROBE-AT with transcriptional fusions of nptII promoter-gfp and hrpA promoter-mCherry, *hrpA* promoter from *Dickeya dadantii* 3937, Ap^R^ | [5] |
| pAT-P*hrpA_Ea_-*P*nptII* | pPROBE-AT with transcriptional fusions of nptII promoter-gfp and hrpA promoter-mCherry, *hrpA* promoter from *Erwinia amylovora1189*, Ap^R^ | This study |
| pCL1920 | Spec^r^, | [6] |
| pP*hrpA-*P*nptII* | pCL1920 with transcriptional fusions of nptII promoter-gfp and hrpA promoter-mCherry, *hrpA* promoter from *Erwinia amylovora1189*, Spec^R^ | This study |
| pP*gapA-*P*nptII* | pCL1920 with transcriptional fusions of nptII promoter-gfp and gapA promoter-mCherry, *gapA* promoter from *Erwinia amylovora1189*, Spec^R^ | This study |
| pP*hrpA-*P*dspE* | pCL1920 with transcriptional fusions of *dspE* promoter-gfp and hrpA promoter-mCherry, *hrpA* and *dspE* promoters from *Erwinia amylovora1189*, Spec^R^ | This study |
| pP*hrpA-*P*hrpJ* | pCL1920 with transcriptional fusions of *hrpJ* promoter-gfp and hrpA promoter-mCherry, *hrpA* and *hrpJ* promoters from *Erwinia amylovora1189*, Spec^R^ | This study |
| Primers |  |  |
| Km_F | CGGGGTACCTGTGTAGGCTGGAGCTGCTTCG | This study |
| Km_R | CGGAATTCCATATGAATATCCTCCTTAGTTCCTATTCC |  |
| pnptII_F | CGGAATTCACTGGGCTATCTGGACAAGG | This study |
| pnptII_R | GTTGGATCCAATCATGCGAAACGATCCTC |  |
| P*hrpL* mut_F | GATGTGGCGGTTGAATGCCGGGTAAAACGGGAGCAATTTTCATCCTGGCGTGTGTAGGCTGGAGCTGCTTCG | This study |
| P*hrpL* mut_R | CCATCGTTGACCGATGTTGATTCAGTTGTTTGCAGGTGAATTTCTGTCATGGCCGCTCTAGAACTAGTGGATCC |  |
| *PhrpA1189_F* | GTTGGATCCCAAGGGGGACATTCTTGTGT | This study |
| *PhrpA1189_R* | GGTGAGCTCGCCGCTCATATTAATCTCTCCA |  |
| *PgapA1189_F* | GTTGGATCCCAACCTTTGCAGAGCTGGTT | This study |
| *PgapA1189_R* | GGTGAGCTCCAGCTTGTTGATAGTGAATAAA |  |
| *PdspE1189_F* | CGGGATCCCCAGCGTTACCAACTCCTGT | This study |
| *PdspE1189_R* | CGGAATTCGACCCGTTGCCCCCAC |  |
| *PhrpJ1189_F* | CGGGATCCCCAGGTCATTTGCTCCAGAT | This study |
| *PhrpJ1189_R* | CGGAATTCCCTGGCGAACCTTCAATGAT |  |
| gfp_R | CGGGGTACCTTTATGCTTCCGGCTCGTAT | This study |
| mCherry_F | TCCGTCGACTTACTTGTACAGCTCGTCCA |  |
| *hrpA* qPCR_F | CAGCAATGGCAGGCATGCAG | [7] |
| *hrpA* qPCR_R | CTGGCCGTCGGTGATTGAGC |  |
| *dspE* qPCR_F | GATGGCGGAGCTGAAATCGTTC | [7] |
| *dspE* qPCR_R | CCTTGCCGGACCGCTTATCATT |  |
| *rplU* qPCR_F | GCGGCAAAATCAAGGCTGAAGTCG | [7] |
| *rplU* qPCR_R | CGGTGGCCAGCCTGCTTACGGTAG |  |

^a^ Cm^R^, Chloramphenicol resistance; Km^R^, Kanamycin resistance; Am^R^, Ampicillin resistance; Spec^R^, Spectinomycin resistance; gfp, green fluorescent protein gene; mCherry, red fluorescent reporter protein gene. Sequence underline indicated restriction enzyme sites.
