## Supplementary figures and images for "Expression of the type III secretion system genes in epiphytic *Erwinia amylovora* cells on apple stigmas benefits endophytic infection at the hypanthium"

### Supplemental Figure 1

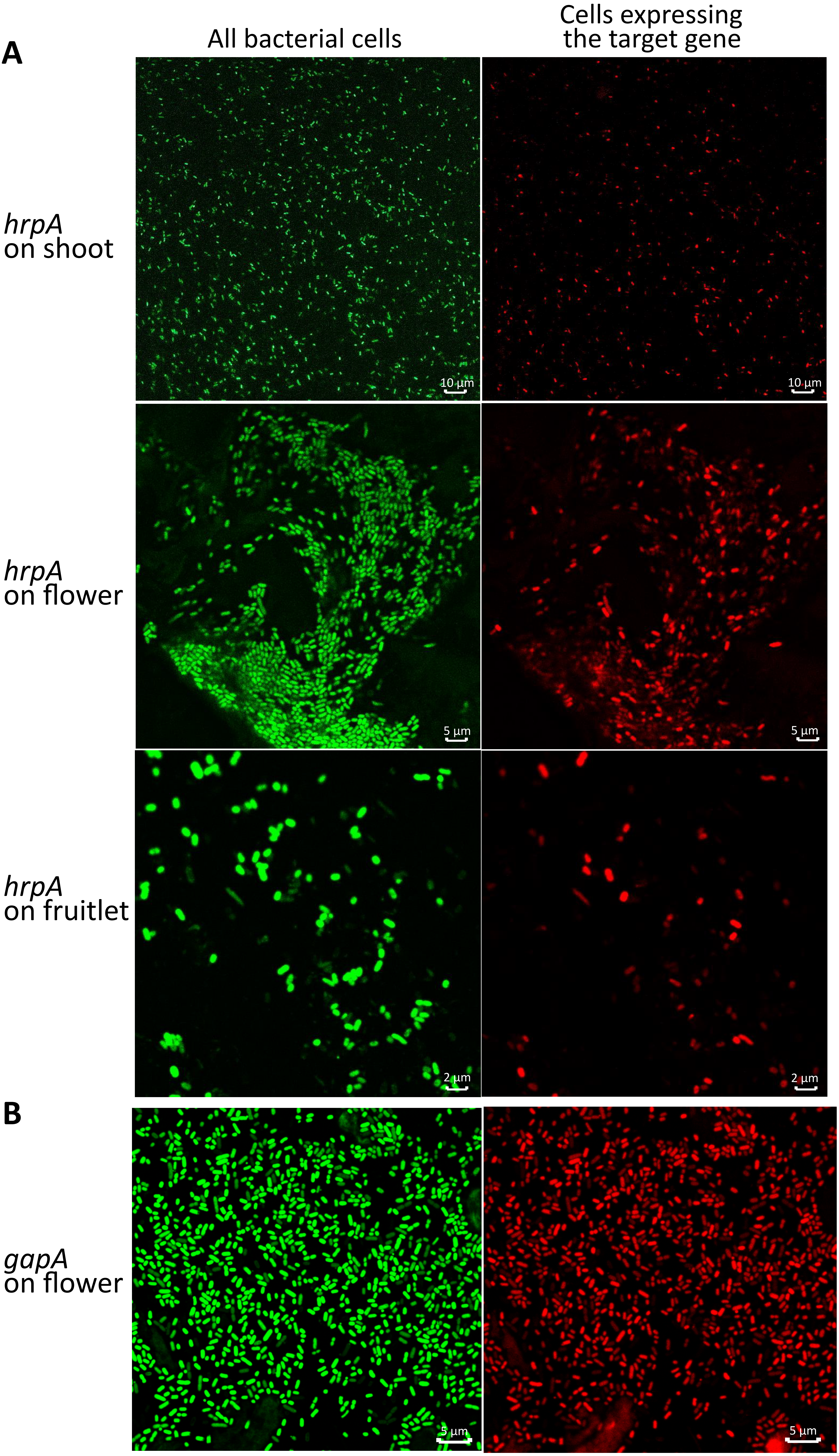

### Supplemental Figure 2

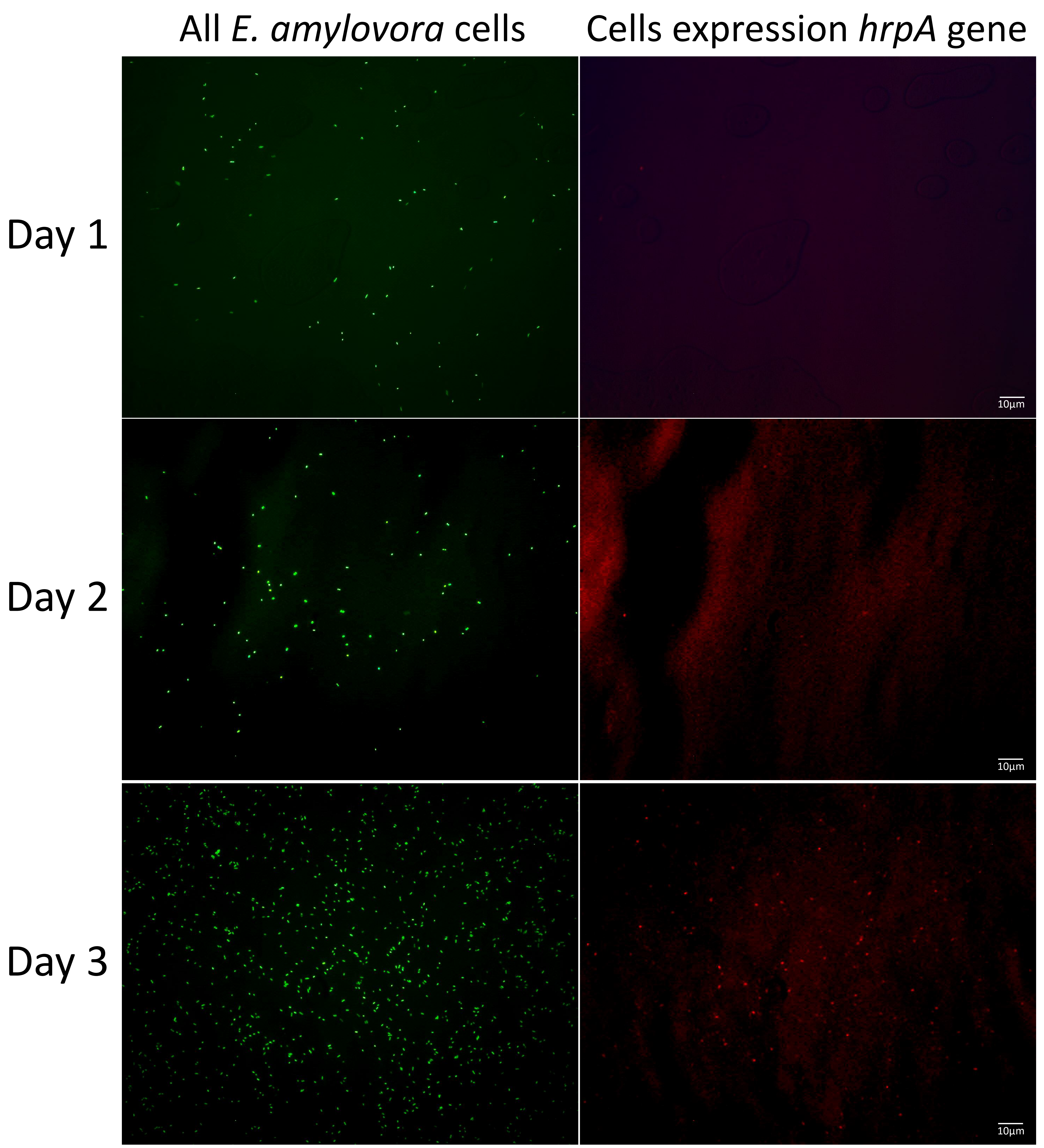

### Supplemental Figure 3

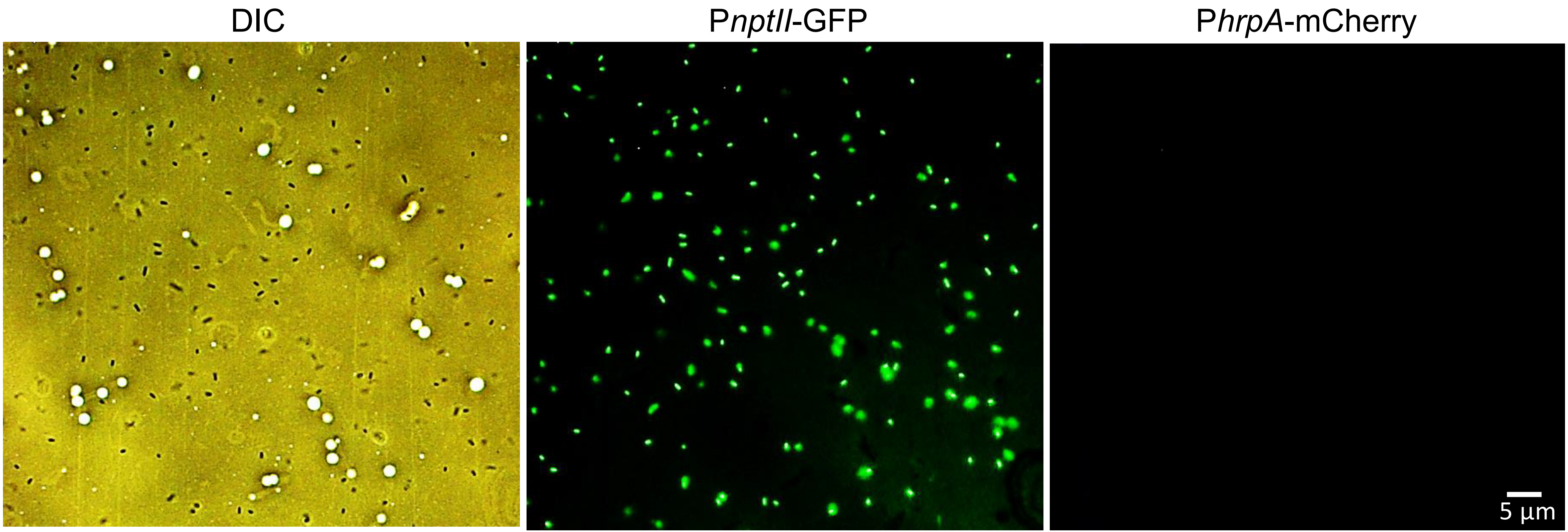
